# Historical records of plant-insect interactions in subarctic Finland

**DOI:** 10.1101/2022.01.25.477384

**Authors:** Leana Zoller, Tiffany M. Knight

## Abstract

Historical ecological records document the diversity and composition of communities decades or centuries ago and provide a valuable benchmark for modern comparisons. Historical datasets on plant-animal interactions allow for modern comparisons that examine the stability of species and interaction networks over long periods of time and in response to anthropogenic change. Here we present a curated dataset of interactions between plants and insects in subarctic Finland, generated from digitizing a historical document from the late 19th century and updating the taxonomy using currently accepted nomenclature. The resulting dataset contains 654 records of plant-insect interactions observed during the years 1895-1900, and includes 498 unique interactions between 86 plant species and 173 insect taxa. Syrphidae, Apidae and Muscidae were the insect families involved in most interactions, and interactions were most observed with the plant species *Angelica archangelica, Salix caprea*, and *Chaerophyllum prescottii*. Interaction data are available as csv-file and provide a valuable resource on plant-insect interactions over 120 years ago in a high latitude ecosystem that is undergoing rapid climate change.

## Background & Summary

The rapid degradation of natural ecosystems in the Anthropocene^1,2^ highlights the increasing need for conservation actions that preserve life-sustaining ecosystem functions and services^3^. Pollination is a vital ecosystem service as most angiosperm plants, including many crops, rely on animal pollination for sexual reproduction^4,5^. There have been recent observations of declines of pollinators and the plants they are associated with^6^, driven by intensive agriculture, pesticides, the spread of invasive species and pathogens, and climate change^7^. It may take decades or centuries for the full effects of these drivers on plant-pollinator interactions to be realized, and short-term studies may therefore underestimate their effects. Currently, our knowledge on temporal and spatial changes in plant-pollinator interactions is limited, as the vast majority of studies documenting plant-pollinator interactions encompass only one or a few years of the present^8^ and come from North America and Western Europe^9^.

One way to bridge this knowledge gap is through the use of historical records, especially from understudied regions (e.g. tropical and arctic regions). Plant-pollinator visitation networks are constructed through observations of insects coming into contact with the reproductive organs of flowers. Historical datasets documenting these field observations provide rare opportunities to examine long-term changes in pollinator communities and the structure of plant-pollinator networks. For example, Burkle and colleagues^10^ reconstructed a plant-bee visitation network from the late 1800s in Illinois (USA) using a historical document^11^. They resampled the study location, and documented that 55% of the bee species were locally extirpated. Remaining species dramatically restructured their interactions, likely due to spatial and temporal mismatches between interacting species caused by habitat fragmentation and climate change. Research from other areas of the world are urgently needed to understand the generality of these results^12^. For example from arctic and subarctic regions, which are experiencing more rapid climate change compared to the global average^13^ and where flies are the most important pollinators^14,15^. Historical datasets from these regions would provide an important benchmark of plant-pollinator interaction structure, enabling many modern research questions in pollination ecology.

Here, we present a digitized dataset on plant-insect interactions in subarctic Finland derived from a historical document. In the years 1895-1900, Frans Silén observed interactions between plants and insects in Kittilä, Finland and published these observations in the naturalist journal *Meddelanden af Societas pro Fauna et Flora Fennica*^16^. Kittilä is located ~120 km north of the Arctic Circle in a boreal biome. Silén’s original publication is written in Swedish language and consists of a list of observations of 86 plant species visited by a total of 187 insect taxa, resulting in 503 unique interactions. Further, date (day, month and year) and verbatim locality of the observation as well as information on sex, behaviour, and insect quantity in categories (e.g. “scarce”, “many”) along with additional field notes and comments are included. Both plant and insect names were validated to match the currently accepted nomenclature and their higher taxonomical classifications were extracted. After validation, the dataset encompasses 173 insect taxa interacting with 86 plant species, resulting in 498 unique interactions.

## Methods

In a first step, Silén’s original records were manually digitized (InteractionData_Silen.csv). Each unique plant-insect interaction per site and date was entered as a new row of data (hereafter referred to as ‘record’). Full verbatim taxonomic species names of plants and pollinators (as originally stated in the historical document), verbatim locality and date (year, month and day) were included. Additional information on insect sex (i.e. m/f), insect behaviour (e.g. nectar sucking) and categorical abundance (e.g. “scarce”, “many”) was available for many records. We included categorical abundance in the original Swedish language and also provided an English translation. Some records in the historic document contained additional comments or field notes and they were also included in the dataset, but only as English translation. In a second step, verbatim taxonomic plant and insect names were updated to currently accepted names (see Technical Validation section) and added to the interaction dataset.

## Data Records

### Available data formats and structure

The interaction dataset and two datasets containing information on the taxonomic validation of plants and insects are formatted as csv-files (InteractionData_Silen.csv, Plants_TaxonomicValidation.csv and Insects_TaxonomicValidation.csv) and are available on the figshare repository. All column names are described in Tables 1–2.

**Table 1.** Description of the columns labels used in the Interaction dataset (InteractionData_Silen.csv).

| column label | column description | example |
| --- | --- | --- |
| verbatimLocality | The original textual description of the place of recording as it appeared in the original record | Kittilä |
| country | The country in which the verbatim locality occurs | Finland |
| day | The integer day of the month on which the event occurred | 29 |
| month | The integer month in which the event occurred | 6 |
| year | The four-digit year in which the event occurred | 1898 |
| eventDate | The date when the event was recorded | 1898-06-29 |
| plantVerbatimIdentification | The unaltered original taxonomic identification of the plant as it appeared in the original record, including uncertainties, etc. | Trientalis europaea L. |
| animalVerbatimIdentification | The unaltered original taxonomic identification of the insect as it appeared in the original record, including uncertainties, etc. | Syrphus luniger Meig. |
| plantScientificName | The full scientific name of the plant at the lowest level taxonomic rank that can be determined | Lysimachia europaea (L.) U.Manns & Anderb. |
| plantFamily | The full scientific name of the family in which the plant taxon is classified | Primulaceae |
| plantGenericName | The genus part of the plantScientificName without authorship | Lysimachia |
| plantSpecificEpithet | The name of the species epithet of the plant in plantScientificName | europaea |
| plantTaxonRank | The taxonomic rank of the most specific name of the plant in the plantScientificName | species |
| animalScientificName | The full scientific name of the animal at the lowest level taxonomic rank that can be determined | *Eupeodes luniger* (Meigen, 1822) |
| animalOrder | The full scientific name of the order in which the animal taxon is classified | Diptera |
| animalFamily | The full scientific name of the family in which the animal taxon is classified | Syrphidae |
| animalGenericName | The genus part of the animalScientificName without authorship | *Eupeodes* |
| animalSpecificEpithet | The name of the species epithet of the animal in animalScientificName | *luniger* |
| animalTaxonRank | The taxonomic tank of the most specific name of the animal in the animalScientificName | species |
| animalSex | The sex of the animal represented in the occurrence (f = female, m = male, m/f = both) | f |
| verbatimAnimalQuantity | A number or enumeration value for the quantity of animals in the language of the original record | talrik |
| animalQuantity | A number or enumeration value for the quantity of animals, translated to English | numerous |
| animalBehavior | A description of the behavior shown by the animal at the time the occurrence was recorded, translated to English | sucking nectar |
| fieldNotes | Notes taken about the Event | Visitor persistently sucking nectar |

**Table 2.**
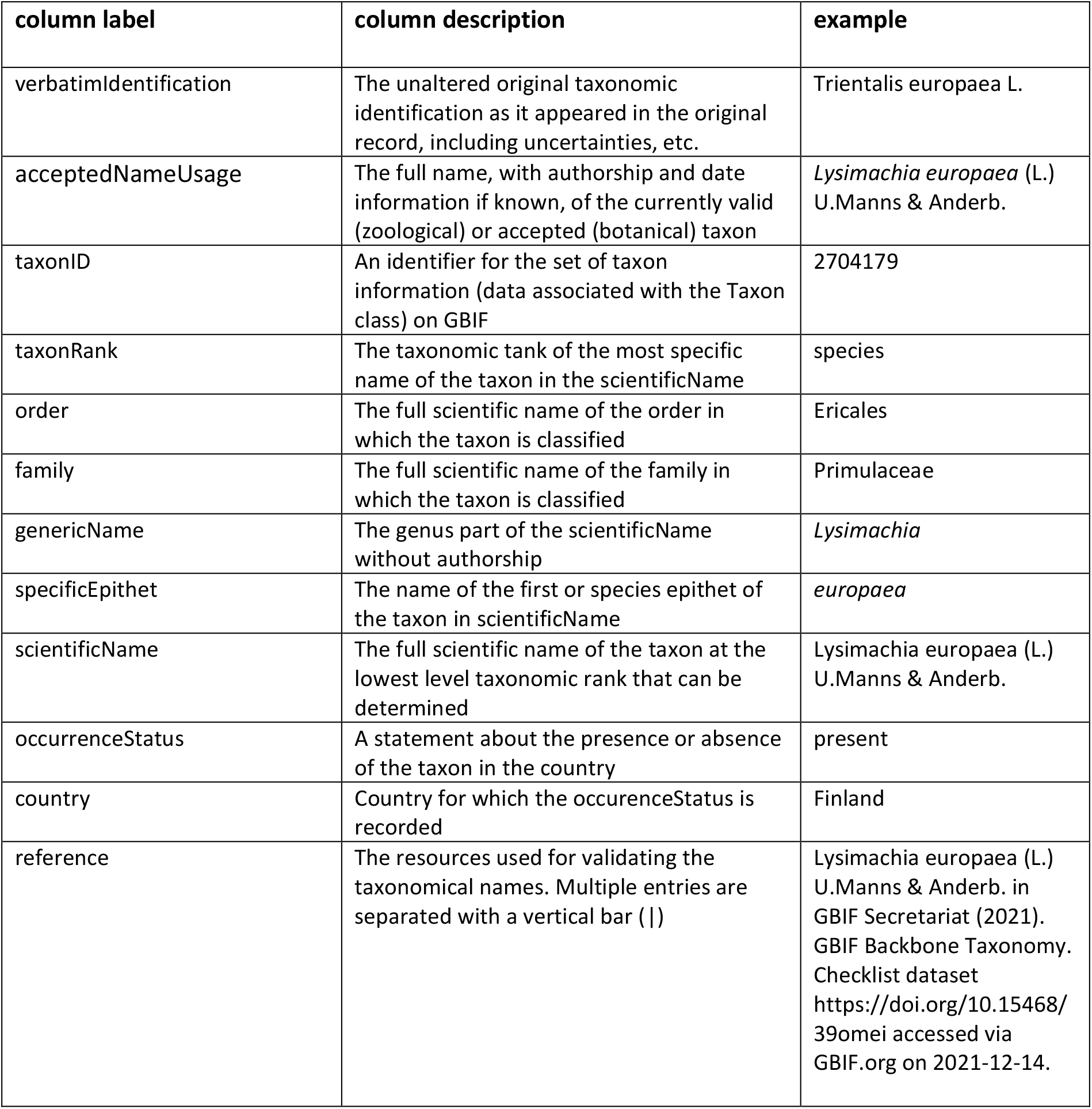
Description of the columns labels used in the Taxonomic validation datasets (Plants_TaxonomicValidation.csv and Insects_TaxonomicValidation.csv.

### Data characterization

In the sections below, we characterize the geographic, taxonomic and temporal coverage of the interaction data.

### Geographic coverage

Records stem from the region around Kittilä, Finnish Lapland (67°39’58.3”N 24°53’25.8”E).

### Taxonomic coverage

Originally, Silén’s data included 654 records of 187 insect taxa visiting 86 plant species, resulting in a total of 503 unique interactions. Of the 187 insect taxa identified by Silén, 164 were resolved to species level (94.95% of records). Among them, three species (6 records) contained information on subspecies. Nineteen taxa were resolved to genus level (4.28% of records) and five taxa were resolved to subfamily, family or superfamily level (0.76% of records). Plant species were all resolved to species level, among them, three species (18 records) contained information on subspecies. After cross-checking taxonomic names, 153 taxa were resolved to species level (94.34% of records), 13 to genus resolution (2.60% of records), six to family level (2.14% of records) and one to order level (0.92% of records). All plant species could be resolved to species level. The recorded insect species belong to four orders (Diptera, Hymenoptera, Lepidoptera and Coleoptera) and include 88 genera in 30 families. The most frequently recorded insect families were Syrphidae, Apidae and Muscidae (Fig. 1) and the most frequently recorded genera were *Bombus, Platycheirus* and *Thricops* (Fig. 2a). Salicacea, Apiaceae and Asteracea were the most frequently recorded plant species, (Fig. 1), and in particular the plant species *Angelica archangelica, Salix caprea*, and *Chaerophyllum prescottii* (Fig. 2b).

**Fig. 1.**
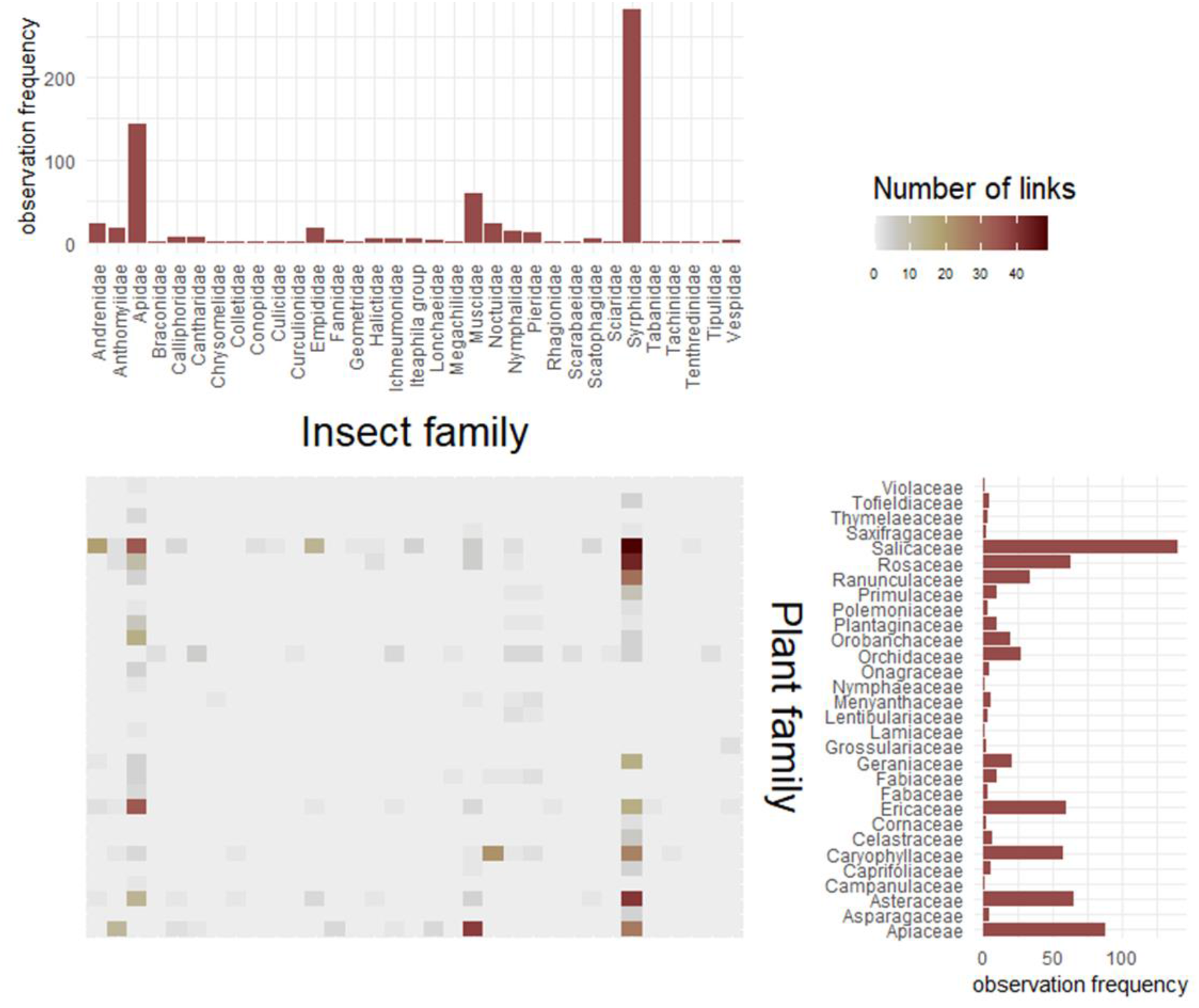
Overview of the number of records (number of times an interaction between a plant species and insect taxa was observed across all sites and dates). This information is summed for each insect family (top) and plant family (right) to allow visualization of the most commonly recorded families and interaction combinations. Six insect records that were identified to a level coarser than family were excluded and information on the categorical quantity of the insects (as stated in the historical source) is not included.

**Fig. 2.**
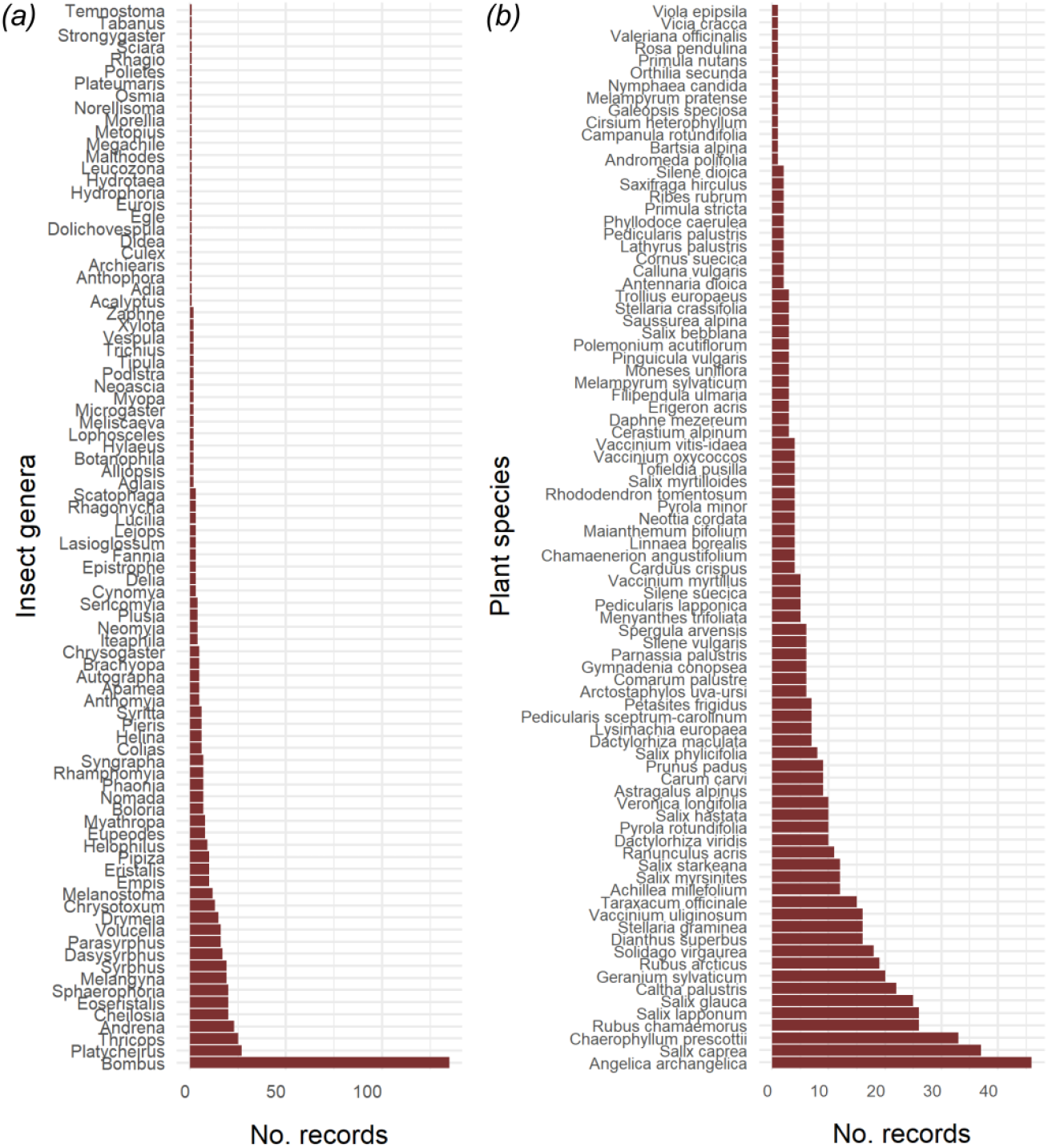
Taxonomic coverage of records. Overview of the number of records in the dataset by (**a**) insect genera and (**b**) plant species. Six insect records that were identified to a level coarser than genus were excluded from the figure. Information on the categorical quantity of the insects is not included in the number of observations.

### Temporal coverage

The records span 126 days between May and August of the years 1895-1900. Six records had information on neither day, month nor year, and 11 records included information on year, but not month or day. The bulk of the records (60.91%) stem from the years 1896 and 1897 and the months June and July (76.3%) (Fig. 3).

**Fig. 3.**
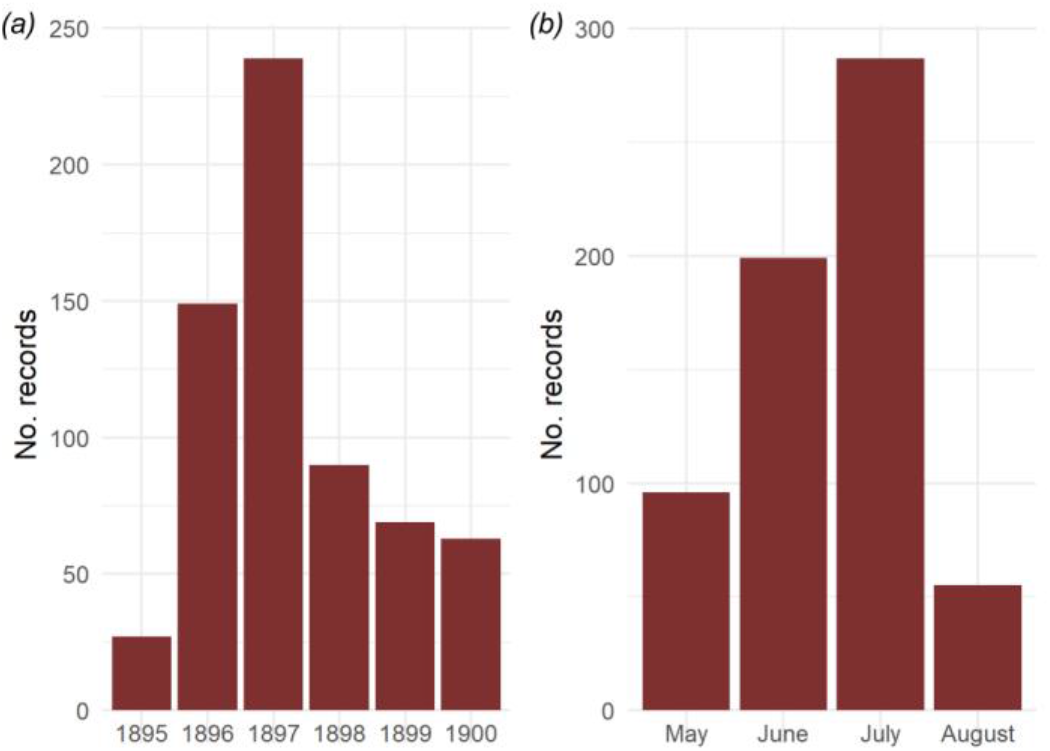
Temporal coverage of records. (**a**) Yearly distribution of plant-insect observations in the dataset and (**b**) monthly distribution of plant-insect observations in the dataset. Seventeen records that did not have information on year and month were excluded from the figure.

## Technical Validation

Each unique verbatim taxonomic name was cross-checked with the GBIF Backbone Taxonomy and Finnish species checklists and, if necessary, the taxonomic name was updated to the currently accepted name (according to the GBIF Backbone taxonomy). Additionally, we extracted information on order, family, and genus of each taxon. When verbatim taxonomic names could not be resolved to a valid taxon using the GBIF Backbone Taxonomy and checklists, we manually researched taxonomic revisions of the verbatim taxa in other databases, publications or checklists. When the verbatim species names could not be resolved to any currently valid species, the next finest available resolution (genus, family or order), was recorded. Further, we verified if the derived species have previously been reported from Finland using the online portal (laji.fi) of the Finnish Biodiversity Information Facility (FinBIF). Verbatim taxonomic names with corresponding updated names, sources for the new names, and information of occurrence in Finland as well as the GBIF identifiers of each taxon are provided for plants and insects in two supplementary data files (Plants_TaxonomicValidation.csv and Insects_TaxonomicValidation.csv).

## Code Availability

No custom code was used to generate the data described in the manuscript. Code used to create summarizing figures is available online (https://github.com/LeanaZ/historic-interactions).

## Acknowledgements

We thank J. Everaars for bringing this historical text to our attention, P. Schnitker for help with data digitization and J. Kahanpää for expert advice on Diptera taxonomy. This research was supported by the Alexander von Humboldt professorship and the Helmholtz Recruitment Initiative, both awarded to TMK, and by the support of iDiv by the German Research Foundation (FZT 118).

## Author contributions

TMK and LZ conceived the ideas and designed the methodology; LZ led the data digitization; LZ led the writing of the manuscript. TMK contributed critically to the drafts and gave her final approval for publication.

